# V-ATPase controls tumor growth and autophagy in a *Drosophila* model of gliomagenesis

**DOI:** 10.1101/2020.07.20.211565

**Authors:** Miriam Formica, Alessandra Storaci, Irene Bertolini, Valentina Vaira, Thomas Vaccari

**Author notes:** These authors contributed equally (Valentina Vaira, Thomas Vaccari). Electronic address.

## Abstract

Glioblastoma (GBM), a very aggressive and incurable tumor, often results from constitutive activation of EGFR (epidermal growth factor receptor) and of PI3K (phosphoinositide 3-kinase). To understand the role of autophagy in the pathogenesis of glial tumors *in vivo*, we used an established *Drosophila melanogaster* model of glioma based on overexpression in larval glial cells of an active human *EGFR* and of the PI3K homolog *Dp110*. Interestingly, the resulting hyperplastic glia expresses high levels of ref(2)P (refractory to Sigma P), the *Drosophila* homolog of p62/SQSTM1. However, cellular clearance of autophagic cargoes appears inhibited upstream of autophagosome formation. Remarkably, downregulation of subunits of the vacuolar-H+-ATPase (V-ATPase) prevents overgrowth, reduces PI3K signaling and restores clearance. Consistent with evidence in flies, neurospheres from patients with high V-ATPase subunit expression show inhibition of autophagy. Altogether, our data suggest that autophagy is repressed during glial tumorigenesis and that V-ATPase could represent a therapeutic target against GBM.

## Introduction

Gliomas, the most common brain malignancy, represent a challenge for therapy because of limited treatment options and of the onset of therapeutic resistance. Among gliomas, GBM is by far the most aggressive and incurable [1], with a 5-year survival rate of only 5% [2]. Even in patients with positive prognostic factors, maximum surgical resection and adjuvant chemoradiotherapy, the overall median survival rate is limited to 14.6 months [3]. The most frequent genetic feature of GBM is mutation of *EGFR*, leading to a constitutively activated form of the receptor in around 40-50% of primary GBMs [4]. The PI3K pathway, which is one of the EGFR effectors can also be mutated in 20% of tumors, contributing to uncontrolled cell growth [5]. Thus, components of the EGFR and PI3K pathways, including the serine/threonine kinase AKT and the mammalian target of rapamycin (mTOR), are widely considered potential targets to develop new GBM treatments in combination with other therapeutics [6], [7].

A recently discovered prognostic feature of GBM is expression of subunits of the V-ATPase proton pump, which are frequently found upregulated in cancer [8], [9]. In fact, in GBM tissue samples and GBM patient-derived neurospheres (NS), increased expression of a subset of V-ATPase subunits positively correlates with GBM aggressiveness and poor patient survival [10], [11]. Interestingly, V-ATPase and the mTOR complex 1 (mTORC1) kinase act with the Transcription factor EB (TFEB) family of lysosomal associated proteins to form a homeostatic circuit that balances catabolic and anabolic processes [12], [13]. When mTORC1 is inactive, TFEB translocates into the nucleus to modulate expression of genes harboring a CLEAR (Coordinated Lysosomal Expression And Regulation) site, thus controlling lysosomal biogenesis and macroautophagy (autophagy hereafter) [14]. However, whether and how V-ATPase regulates tumor growth in genetic models of glioma development is not known.

Cell type-specific regulation, genetic alterations, tumor staging or treatment most likely determine the exact role of autophagy in tumorigenesis [15]–[17]. For instance, during tumor initiation autophagy has been shown to play a tumor-suppressive role. However, once the tumor is established, autophagy can instead positively impact tumor survival by increasing metabolic activity in support of cell proliferation or survival to hypoxia [18]. Importantly, autophagy appears to play different roles in cancer stem cells, compared to differentiated cells, and provides resistance to chemotherapy [19]–[22]. In GBM, treatment with the drug Temozolomide (TMZ) has been reported to trigger autophagy [23], [24] and combination therapy with the V-ATPase inhibitor Bafilomycin A1 (BafA1) increases glioma cell death [10], [25]. Despite this, the role of autophagy in gliomagenesis remains largely underexplored.

In this study, we used *Drosophila melanogaster* as an *in vivo* model to define the role of V-ATPase and autophagy during glioma development. *Drosophila* encodes single homologues of most genes altered in GBM, all displaying high degrees of functional conservation with mammals [26]. Our data indicate that autophagy is repressed both *in vivo* and in patient-derived NS and that V-ATPase is a limiting factor for growth and autophagy inhibition.

## Results and discussion

As previously demonstrated [26], [27], co-expression in *Drosophila* larval glial cells of the constitutively active form of PI3K *Dp110-CAAX* and of human *EGFR ΔhEGFR*, under the control of *repo-Gal4*, a *P(Gal4)* insertion in the glial-specific *reverse polarity* (*repo*) locus, promotes excess cell growth and hypertrophy of the optic lobes of the central nervous system (CNS) **(Fig. S1A-A’’’)**. In control brains, glial cells constitute 10% of the CNS cells, while the rest is mostly formed by neurons and neural progenitors **(Fig. 1A;** [28]). Upon co-expression of *Dp110-CAAX* and *ΔhEGFR*, cells of glial origin make up 70% of the recovered CNS cells **(Fig. 1A)**. Consistent with this, *repo* transcription is upregulated compared to controls, while expression of the neuronal marker *embryonic lethal abnormal vision* (*elav*) is strongly downregulated (**Fig. 1B-C**). Alongside, the corresponding Repo and Elav proteins are similarly deregulated (**Fig. 1D**). Morphologically, larvae carrying gliomas show an extremely altered CNS arrangement, with neurons located in small clusters surrounded by hyperplastic glia **(Fig. S1B-B’)**, and fail to wander or pupariate, eventually dying at third instar **(Fig. S1C-C’’)**. These data confirm that the cell growth aspects of gliomagenesis can be recapitulated in *Drosophila* and extend the description of such *in vivo* genetic model.

**Figure 1.**
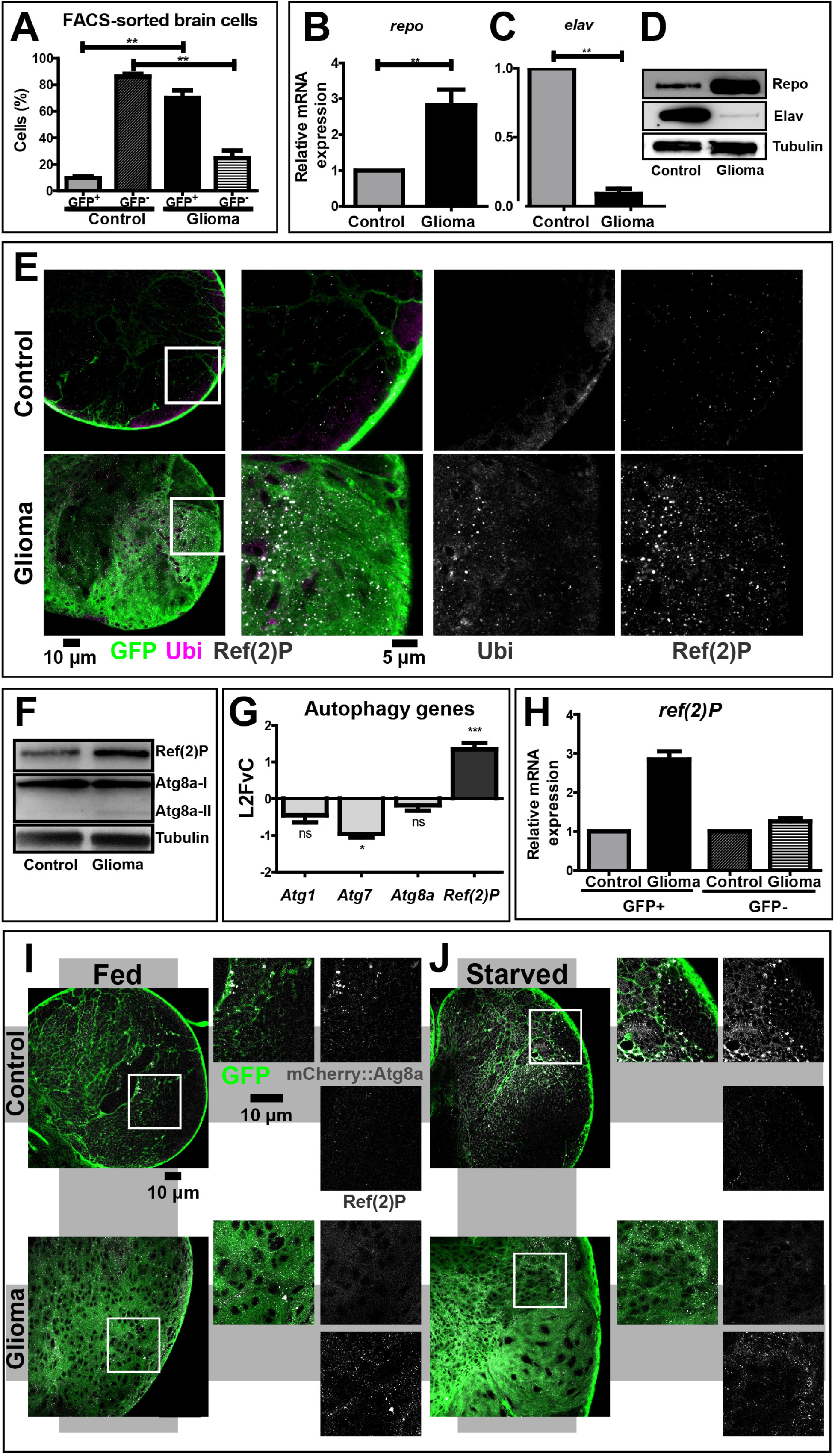
The autophagy-lysosomal pathway is inhibited during glial overgrowth induced by expression of *Dp110* and *ΔhEGFR*. (A) Larval brain cells were separated by FACS. In controls, glial cells (GFP+) represent 10% of the total brain population, while, in gliomas, glial cells are up to 70%. Accordingly, glia overgrowth involves a strong reduction of neurons (GFP−) which are heavily decreased compared to control brains. Data represent the mean ± S.D. of n≥3 independent experiments and *P-values* are determined by Kruskal Wallis test with Dunn’s Multiple Comparison. (B-C) mRNA expression of *repo* (glia) and *elav* (neurons). *repo* levels are strongly increased in gliomas compared to controls. Conversely, *elav* levels are heavily suppressed in gliomas samples. *RpL32* is used as a housekeeping control. Data represent the mean ± S.D. of n≥3 independent experiments and *P-values* are determined by Mann-Whitney test. (D) Western blot showing Repo and Elav protein levels. Repo levels confirm a strong expansion of glial tissue in gliomas, while Elav protein levels are strongly decreased in tumor brains. Tubulin is used as a loading control. (E) Single medial confocal sections third instar larval brains. High magnification insets are shown as merge and separate channels. Glial cell membranes (marked with anti-GFP), ubiquitin (marked by anti- Ubiquitin FK2), Ref(2)P are pseudo-colored as indicated. Notice the increased signal of both markers in gliomas compared to controls. Ubiquitin mostly colocalize with Ref(2)P. (F) Western blot showing expression of the autophagy markers Ref(2)P and Atg8a I-II. In gliomas, ref(2)P protein levels are increased compared to controls, albeit Atg8a levels are only slightly changed. Tubulin is used as a loading control. (G) mRNA levels showing expression autophagy genes. *ref(2)P* mRNA expression in glioma samples is strongly upregulated, while mRNA level of *Atg1, Atg7 and Atg8a* are comparable to controls. Data are expressed as fold increase relative to control brains (L2FvC). Data represent the mean ± S.D. of n≥3 independent experiments and *P-values* are determined by Mann-Whitney test. (H) *ref(2)P* expression levels in GFP+ and GFP-brains cells separated by FACS. *ref(2)P* levels are increased GFP+ cells that belong to tumor tissue, but not in GFP-neural cells. Normalization on GFP+ cells of control brains. *RpL32* is used as housekeeping. Data represent the mean ± S.D. of n≥3 independent experiments. (I-J) Single medial confocal sections of third instar larval brains reared under fed and starved conditions. Glial cell membranes are marked by GFP, Atg8a is detected by mCherry (mCherry::Atg8a). Note the strong increase of mCherry::Atg8a signal upon starvation in controls but not in gliomas. In contrast, upon starvation of gliomas ref(2)P is strongly accumulated.

To characterize the role of the autophagy-lysosomal pathway in gliomagenesis *in vivo*, we evaluated the presence of accumulations of ubiquitin and of the autophagy-specific cargo adapter Ref(2)P (refractory to Sigma P; p62/SQSTM1 in mammals) in larval brains. Compared to control glia, in which very little signal of either marker is detected, both proteins strongly colocalize in puncta within glial cells of larvae carrying gliomas **(Fig. 1E)**. This result suggests that during gliomagenesis the autophagic process could be either impaired or heavily induced. To discriminate, we assessed the autophagic flux by monitoring the expression of Ref(2)P and of Atg8a (Autophagy-related protein 8a; LC3 in mammals). We confirmed that Ref(2)P accumulates in glioma CNS extracts, while Atg8a levels are only slightly increased, when compared to control extracts **(Fig. 1F**). In addition, we found that transcription of *Atg8a* and other core autophagy genes, such as *Atg1* and *Atg7* is mostly unchanged relative to controls, while that of *ref(2)P* is upregulated **(Fig. 1G)**. Then, we sorted cells to reveal that *ref(2)P* expression is increased exclusively in GFP+ glial cells belonging to tumor brains **(Fig. 1H)**. We next used starvation, a known stimulus triggering autophagy, to evaluate whether such pathway could be induced during gliomagenesis. Consistent with basal levels of constitutive autophagy, mCherry::Atg8a a sub-cellular marker of autolysosome formation can be detected in control glial tissue but not in glioma cells under fed condition **(Fig. 1I)**. Upon starvation, the mCherry::Atg8a signal is strongly increased in control samples, indicating induction of autolysosome formation by nutrient deprivation **(Fig. 1J)**. In stark contrast, the CNS of larvae carrying gliomas shows no appreciable increment of mCherry::Atg8a expression **(Fig. 1J)**. Western blot to detect ref(2)P confirmed that its accumulation decreases during starvation in control, while it remains unchanged in glioma samples **(Fig. S1D)**, indicating the inability to clear autophagic cargoes by autophagy.

To test whether autophagy is also inhibited in patient-derived GBM neurospheres (NS), we examined the morphology of degradative organelles by electron microscopy (EM). We briefly treated NS with the V-ATPase inhibitor BafA1 that blocks fusion of autophagosomes to lysosomes [29], and quantified the number of autophagosomes, which is expected to accumulate upon treatment only in spheres with active autophagy. We found that in NS with a low level of V-ATPase G1 subunit expression (V1G1^Low^NS), BafA1 administration leads to major accumulation of autophagic structures (**Fig. 2A**). In contrast, in NS from patients with elevated expression of V-ATPase G1 (V1G1^High^NS), drug treatment does not significantly change the number of autophagic structures, which is comparable to untreated controls (**Fig. 2A**). Quantification confirms that addition of BafA1 drives the accumulation of abnormal autophagic structures only in V1G1^Low^NS (**Fig. 2A’**). Overall, these findings suggest that in both fly gliomas and patient-derived NS with high V1G expression, autophagy is inhibited upstream of autophagosome formation.

**Figure 2.**
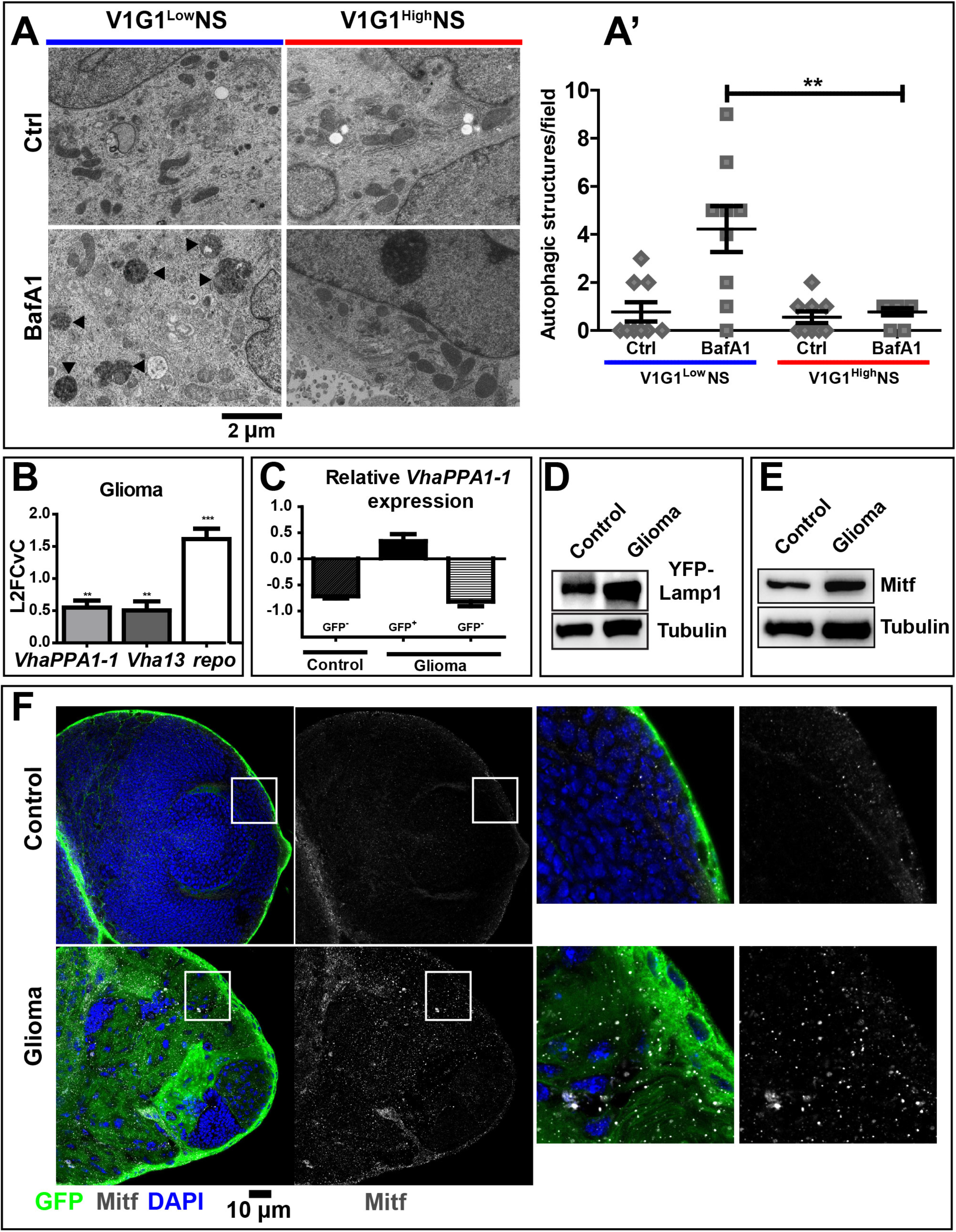
Autophagy in NS and characterization of the lysosomal compartment during gliomagenesis. (A) Representative EM images of NS treated with vehicle (Ctrl) BafA1. V1G1^High^ and V1G1^Low^ NS reveal a net difference in accumulation of autophagic organelles (arrowheads) upon treatment. (A’) Quantification of autophagic structures confirms that BafA1 causes accumulation of aberrant organelles only in V1G1^Low^ NS. Data represent the mean ± S.D. of n≥2 independent experiments, *P-value* is obtained by ANOVA with Bonferroni’s multiple comparison post-test. (B) qPCR analysis of *VhaPPA1-1* and *Vha13* and *repo* expression in fly glioma cells. mRNA expression levels confirm the upregulation of the V-ATPase in fly gliomas. Data are expressed as L2FvC. Bars, mean ± S.D. of n≥3 independent experiments. (C) *VhaPPA1-1* mRNA levels in disaggregated cells are downregulated in GFP-cells, conversely, this subunit is upregulated in GFP+ cells that belong to the tumor. Normalization is relative to GFP+ cells of control brains. Data represent the mean ± S.D. of n≥3 independent experiments. (D) Western blot showing expression of YFP::Lamp1 which localizes to lysosomes. In gliomas, Lamp1 protein levels are increased compared to controls. Tubulin is used as a loading control. (E) Western blot showing expression of Mitf reveal increased protein levels in gliomas. Tubulin is used as a loading control. (F) Single medial confocal sections of third instar larval brains. Nuclei were stained with DAPI, glial cell membranes with anti-GFP. Mitf is heavily accumulated in glioma compared to control optic lobes. Mitf accumulation can be better appreciated in higher magnifications of insets. Notice that in gliomas, Mitf is almost exclusively in the cytoplasm, the transcription factor is seldom associated with nuclei (insets).

Since autophagy and V-ATPase are part of lysosomal nutrient-regulation circuits, we next assessed the lysosome abundance and function in cells undergoing gliomagenesis in *Drosophila*. Interestingly, expression of some subunits of the proton pump, such as *Vha13*, encoding the *Drosophila* subunit V1G, and *VhaPPA1-1,* encoding the *Drosophila* subunit V0C, is increased in larvae carrying gliomas, when compared to controls **(Fig. 2B)**, echoing the elevated expression observed in aggressive GBM NS. FACS analysis indicates that expression of such V-ATPase subunits is higher in glial cell than in other CNS cell types **(Fig. 2C**, **Fig.S2A)**, as suggested by previous evidence on glial functions [28], [30]. In addition, protein and mRNA expression of the lysosomal-associated membrane protein 1 (*Lamp1*) marker is also elevated **(Fig. 2D**, **Fig. S2B)**. Similarly, expression of *Mitf* (microphthalmia-associated transcription factor), the unique fly TFEB homolog [31]–[33], is increased in gliomas compared to controls **(Fig. 2E)**. However, immunofluorescence analysis revealed that Mitf is not appreciably present in the nuclei of glioma cells, compared to those of control glia, suggesting that its increased levels do not correlate with increased activity **(Fig. 2F**). In agreement with this observation, downregulation of *Mitf* during gliomagenesis only minimally affect tumor growth **(Fig. S2C-E)**. Finally, the degradative ability of lysosomes, measured by DQ-bovine serum albumin (BSA) uptake (see Material and methods) is preserved, if not increased, in the CNS of larvae carrying gliomas **(Fig. S2F)**. These data suggest during fly gliomagenesis the lysosomal compartment of glial cell is moderately expanded and active, while TFEB is mostly inactive and not strongly contributing to tumor growth.

Spurred by the elevated expression of certain V-ATPase subunits in fly gliomas and in patient-derived NS, we next determined whether hyperplastic glial growth is sensitive to downregulation of V-ATPase subunits. In contrast to *Mitf* downregulation, glial-specific downregulation of the V-ATPase subunit *VhaPPA1-1* in the context of *Drosophila* gliomagenesis (gliomas>*VhaPPA1-1RNAi*) strongly prevents glial cell overgrowth (**Fig. 3A; S3A**; [11]). In fact, glia-associated GFP expression as well as CNS size are significantly decreased by *VhaPPA1-1* subunit downregulation **(Fig. 3A’-A”)**. In addition to reduced glial overgrowth, we also find that Elav protein levels are increased in gliomas>*VhaPPA1-1RNAi*, when compared to glioma CNSs, indicating a partial reversion of the neuronal loss induced by gliomagenesis **(Fig. S3B)**. Despite this, lethality and alteration of CNS architecture are not rescued by V-ATPase subunit downregulation (**Fig. S3C**), suggesting that not all aspect of gliomagenesis are reverted by *VhaPPA1-1* subunit downregulation. To investigate whether glial downregulation of *VhaPPA1-1* restricts growth by causing cell death, we evaluated tissue expression of the apoptotic marker cleaved caspase-3. Unexpectedly, we found that in controls>*VhaPPA1-1*^*RNAi*^ apoptosis is strongly induced in both glia and neurons, consistent with the possibility that V-ATPase is essential autonomously and non-autonomously for CNS health **(Fig. S3C’)**. However, this is not the case in larvae carrying gliomas **(Fig. S3 C’)**, which we previously found to contain higher glial V-ATPase subunit expression than in healthy larvae **(Fig. 2; S2)**. This result reveals that VhaPPA1-1 activity is essential for survival of otherwise wild-type glial cells, but not for the survival of overgrowing glia.

**Figure 3.**
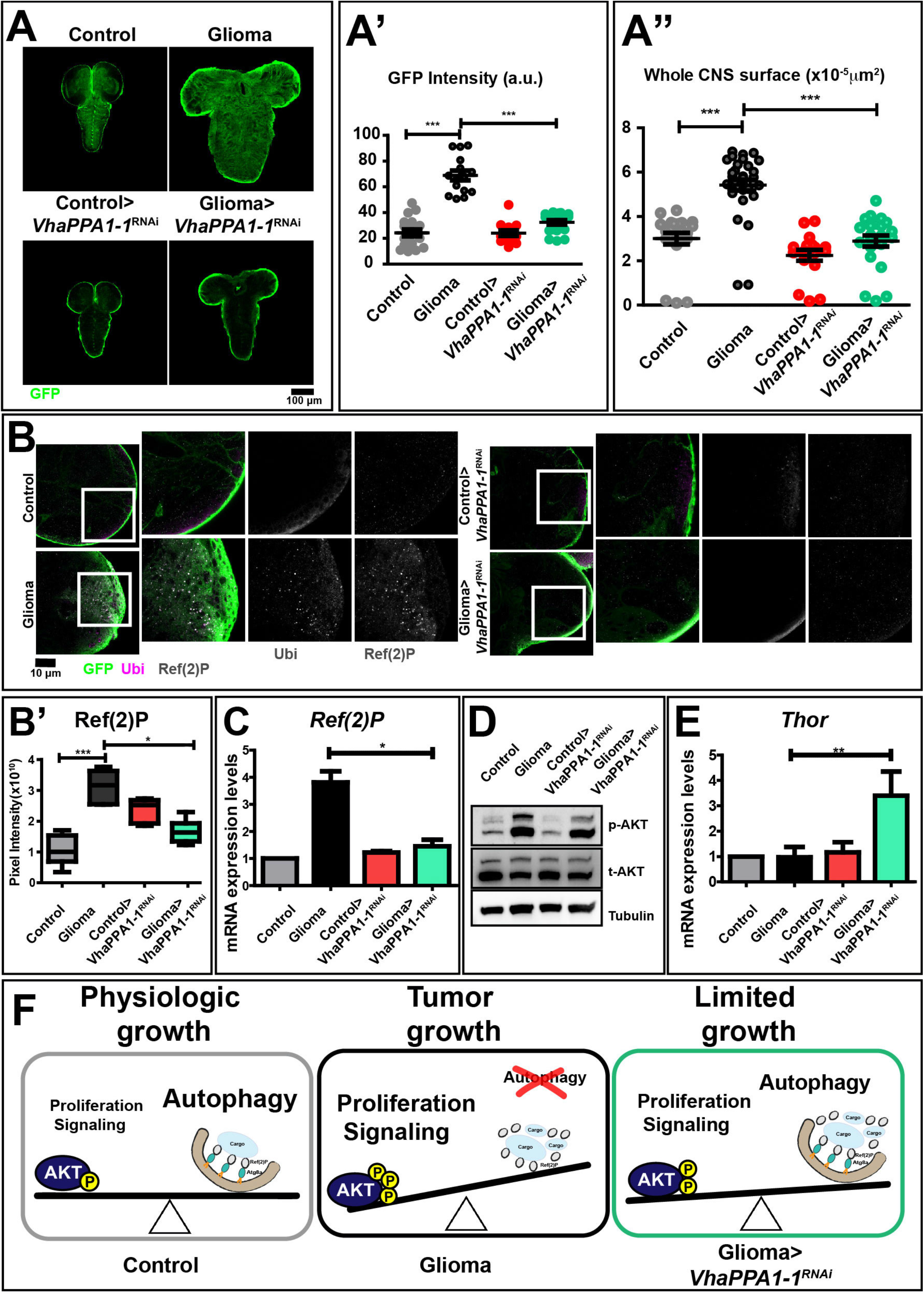
Downregulation of *VhaPPA1-1* restores normal growth and autophagy in cells subjected to gliomagenesis. (A) Single medial confocal sections of a whole CNS from third instar larvae. Dorsal view, anterior up. The excess growth of the glia, observed in gliomas, is strongly reduce to control levels in gliomas>*VhaPPA1-1^RNAi^*. (A’-A’’) Quantification of GFP intensity (A’) and whole CNS surface (A”) confirm that the expansion of glial tissue observed in gliomas is prevented by *VhaPPA1-1* subunit downregulation. Each genotype analysed consists of n≥ 10 CNSa. Mean ± S.D. are shown, and *P*-values are determined by Kruskal Wallis test with Dunn’s Multiple Comparison. (B) Single medial confocal sections of third instar larval CNSs immunostained as indicated. The increased ubiquitin (Ubi) and ref(2)P signal observed in gliomas, is absent in gliomas>*VhaPPA1-1*^*RNA*i^. See also insets on the right of each panel. (B’) Quantification of ref(2)P signal in CNSs. The accumulation of ref(2)P in gliomas is decreased in gliomas in which *VhaPPA1-1* is downregulated. Means ± S.D. are shown, *P*-*values* are determined by One-way ANOVA, Kruskal Wallis test with Dunn’s Multiple Comparison. (C) *ref(2)P* mRNA levels by qPCR. Gliomas>*VhaPPA1-1RNAi* show levels comparable to that of controls. Data represent the mean ± S.D. of n≥3 independent experiments and *P*-*value* is obtained by Kruskal Wallis test with Dunn’s Multiple Comparison. (D) Western blot showing AKT expression levels. The increased p-AKT levels observed in gliomas are decreased upon *VhaPPA1-1* downregulation. Total (t-AKT) AKT and Tubulin levels are used as control. (E) *Thor* mRNA levels by qPCR are upregulated in gliomas>*VhaPPA1-1RNAi*. Expression levels are relative to control brains. Data represent the mean ± S.D. of n≥3 independent experiments, *P*-*value* was obtained by One-way analysis of variance, Bonferroni’s Multiple Comparison Test. (F) A model for V-ATPase function in *Drosophila* larval gliomas. The physiologic balance between anabolic and catabolic processes governing normal cell growth (Physiologic growth) is heavily compromised in gliomas. Indeed, in tumor brains, growth is enhanced while catabolism is impaired (Tumor growth). Downregulation of *VhaPPA1-1* restores the equilibrium controlling nutrient metabolism, ultimately derepressing autophagy decreasing tumor growth (Limited growth).

To uncover the mechanism that underlies the dependency of glioma growth on V-ATPase, we evaluated ubiquitin and Ref(2)P accumulation within tumor tissue upon modulation of *VhaPPA1-1* expression. In sheer contrast to the strong accumulation observed in glioma tissue, neither markers are found as puncta in tumor tissue depleted of *VhaPPA1-1* **(Fig. 3B; quantification in 3B’)**. Notably, the upregulation of *ref(2)P* at the mRNA and protein level observed in gliomas is reverted upon V-ATPase subunit downregulation **(Fig. 3C)**, as are the protein levels detected upon starvation **(Fig. S3D)**. Also, Mitf accumulation in glioma CNSs is prevented by downregulation of *VhaPPA1-1* **(Fig. S3E-F)**. Finally, lysosomal activity is not altered following *VhaPPA1-1* downregulation, while the partial expansion of the lysosomal compartment observed in glioma CNSs is reverted (**Fig. S3G**), indicating that V-ATPase downmodulation normalizes the alterations of catabolism associated to fly gliomagenesis.

To determine how *VhaPPA1-1* downregulation prevents inhibition of autophagy, we evaluated the activity of growth signaling by detecting phosphorylation of AKT (p-AKT). As expected, we found a sharp increase in p-AKT levels in glioma samples **(Fig. 3D)**. Importantly, such increase is blunted by downregulation of *VhaPPA1-1* **(Fig. 3D)**. In addition, expression of *Thor*, a translation inhibitor that is downregulated by AKT [34], [35], is elevated in gliomas>*VhaPPA1-1*^*RNAi*^, when compared to controls **(Fig. 3E)**, consistent with a reduction of anabolism. Overall, the data concerning *VhaPPA1-1* down-modulation during gliomagenesis suggest that reduction of V-ATPase activity might restore catabolism operated by the autophagy-lysosomal pathway and restrain activation of growth pathways promoted by excess EGFR and PI3K signaling.

*Drosophila* models of tumorigenesis have so far shown that autophagy promotes tumor growth in cancer stem cells in the ovary [36], as well as in *Ras*-, but not in JNK-or Notch-induced tumors in imaginal discs [17], [37]. This study explores for the first time the genetic control of autophagy in a dual oncogene model of gliomagenesis in the context of the V-ATPase/TFEB axis.

Compared to nutrient sensing in non-tumor cells (**Fig.3 G**; Physiologic growth), our data reveal that ectopic activation of PI3K signaling might fuel cell growth, not only by increasing anabolism, but also by decreasing catabolism associated to autophagy activation. Whether this is the case, or whether inhibition of autophagy represents merely a side effect of oncogenic proliferative signaling remains to be determined. Persistent growth signaling might also conflict with lysosomal sensing of available nutrients and/or with changes in nutrient demand experienced by tumor cells, leading to the lysosomal compartment anomalies that we have observed (**Fig.3 G**; Tumor growth).

How could inhibition of V-ATPase prevent glial overgrowth? *ΔhEGFR* and *Dp110-CAAX*-mediated activation of the PI3K/AKT/dTOR pathway could prevent TFEB-mediated transcriptional regulation of genes related to the lysosomal-autophagic pathway, and/or it could directly inhibit Atg1 activity. As recently reported, AKT could also repress TFEB activity in a mTORC1-independent manner [38]. Since we observe that *VhaPPA1-1* downregulation not only decreases AKT activity, but it also results in up-regulation of *Thor*, which is known to prevent anabolic processes such as translation, our work establishes V-ATPase as a key node to control the equilibrium between anabolic and catabolic cellular processes in glial tumors. Downregulation of V-ATPase subunit, possibly by limiting V-ATPase activity and/or lysosomal nutrient sensing, reduces activation of the PI3K pathway and associated promotion of protein biosynthesis, as well as normalizes autophagy and the lysosomal compartment, ultimately leading to a sharp decrease in tumor growth (**Fig.3 G**; Limited growth).

Our fly model recaptures aspects of human glioblastoma, including the following evidence obtained with mammalian models and patient samples: Elevated expression of V-ATPase subunits [10], [39]; Induction of autophagy by AKT inhibitors in glioma cells [40]; Proliferation arrest induced by PI3K/mTOR dual inhibitors [41]; Reduction of tumor growth and induction of autophagy by downregulation of the PI3K-AKT-mTOR pathway [42]. Thus, we predict that future study of fly gliomas could provide a framework to uncover new genetic vulnerabilities in GBM. In addition, our model might provide a valuable entry point to test *in vivo* the efficacy of inducers of autophagy and of other modulators of the V-ATPase/TFEB axis as growth inhibitors. Finally, because standard treatment of gliomas with temozolomide induces autophagy and the combination of TMZ with BafA1 enhances cell death in glioma cells [23], [43], future evidence obtained with the fly model could direct us to an informed development of TMZ-based combination therapies.

Despite the evolutionary distance, the ability to model many of the main alterations observed in GBM, as well as the possibility to genetically and pharmacologically interrogate the model, might prove an advantage over monogenic mammalian models. A case in point is that of a murine RAS-only model, which reported an increase of autophagy during gliomagenesis [22]. However, fly gliomas might not recapture complex aspects of GBM, such as the differences observed between glioma stem cells and other glioma cells in terms of regulation of autophagy [21]. Interestingly, an alternative genetic model of gliomagenesis in flies is exists [44] and it could be used to verify outcomes of future experiments. Of note, accumulation of p62 in absence of autophagosome formation has also been observed in mice lacking Atg7, which have been reported to develop spontaneous liver tumors. Interestingly, in such background p62 contributes to tumor progression [45]. Thus, it would be interesting to assess the role of p62 and uncleared cargoes to promoting gliomagenesis, as well as the effect of autophagy modulators during tumorigenesis in highly nutrient-sensitive tissues.

## Methods

### *Drosophila* husbandry

Fly strains were kept and raised into vials containing standard yeast-cornmeal fly food medium. All crosses were performed at 25°C. *Drosophila* lines used in this study were provided by the Bloomington *Drosophila* Stock Center (BDSC, Bloomington, Indiana), the Vienna *Drosophila* Resource Center (VDRC, Vienna, Austria) and by the *Drosophila* Genomics and Genetic Resources (DGGR, Kyoto, Japan). *repo-Gal4, UAS-CD8GFP, UAS-Dp110-CAAX, UAS-ΔhEGFR* were kindly provided by Renee Read (Emory University School of Medicine, Atlanta, Georgia); *mCherry::Atg8a* provided by Gabor Juhasz (Eotvos Lorand University, Budapest, Hungary); *UAS-VhaPPA1-1 RNAi* (47188, VDRC) and *YFP::Lamp1* ([46], DGGR). To induce starvation, larvae were washed in PBS 1X to remove food residues and left for 4 hours on a Petri dish containing sucrose 20% diluted in PBS 1X. After starvation, larvae were dissected to isolate brains for the subsequent analysis. To monitor lysosomal degradation *in vivo*, larvare were treated with DQ-BSA (Sigma) for 6 hrs. All genotypes of the experiments are listed in Supplementary Table 1.

### Immunostaining

Larval brains were fixed using 4% PFA. Tissues were permeabilized with 1X PBS, 1% Triton X-100. Samples were blocked for 30 minutes in 4% BSA diluted in PBST at room temperature. Primary antibody was incubated overnight at 4°C. The secondary antibody was incubated at room temperature for 2 hrs. Primary antibodies against the following antigens were used: Chicken anti-GFP 1:1000 (ab13970, Abcam), mouse anti-FK2 1:250 (BML-PW8810, Enzo), rabbit anti-ref(2)P 1:1000 (a gift from Tor Erik Rusten, Oslo University, Oslo, Norway), rat anti-RFP 1:1000 (5F8, Chromotek), rabbit anti-Mitf 1:200 (developed by our group [33]). Secondary antibodies used were Alexa conjugated secondary antibodies 1:400 (Invitrogen). Samples were mounted on slides using glycerol 70%. Confocal acquisitions were performed using Leica SP2 microscope (Heidelberg, Germany) with ×40/NA 1.25 or ×63/NA 1.4 oil lenses or A1R confocal microscope (Nikon). Measurements and fluorescence evaluation were carried out through the ImageJ Software (National Institutes of Health, Bethesda, USA), images were assembled with Adobe Illustrator.

### qPCR analysis

Larval brains were collected and homogenized using pastels. RNA extraction was performed using RNeasy Mini Kit (Qiagen, 74104). The concentration of extracted RNA was measured using the NanoDrop 1000 Spectrophotometer. Complementary DNA (cDNA) was synthesized from RNA through reverse transcription, according to the SuperScript^®^ VILO™ cDNA Synthesis Kit (Invitrogen, 11754050). RT-PCR was carried out on the ABI/Prism 7900 HT Sequence Detector System (Applied Biosystems, Carlsbad, CA, USA). Samples were tested by Real-time PCR using the following primers that were designed from Universal Probe Library Roche: *Atg8a F_5’CATGGGCTCCCTGTACCA3’, Atg8a R_5’CTCATCGGAGTAGGCAATGT3’; Atg1 F_5’TTTTACGCTGCCCGAACT3’, Atg1 R_5’GCTCCTGTGTCCAGCAGACT3’; Elav F_5’CCGGCAGCACCAGTAAGA3’, elav R_5’CCAGTTGGGGATTGAGGAA3’; ref(2)P F_5’AGACAGAGCCCCTGAATCCT3’, ref(2)P R_5’GGCGTCTTTCCTGCTCTGT3’; Repo F_5’GCATCAAGAAGAAGAAGACGAGA3’, repo R_ 5’GTTCAAAGGCACGCTCCA3’; VhaPPA1-1 F_5’ATCTTCGGTTCGGCCATC3’, VhaPPA1-1 R_ 5’ATAATGGAGTGGCGAAGGAC3’; Thor F_5’CCAGATGCCCGAGGTGTA3’, Thor R_ 5’AGCCCGCTCGTAGATAAGTTT3’.* Amplicon expression in each sample was normalized to *RpL32* mRNA content. The reactions were performed by the Cogentech qPCR service facility (Milan, Italy).

### Western Blot

*Drosophila* larval brains were homogenized with pestles in RIPA buffer plus the addition of proteinase inhibitors 1:200 (539134, Calbiochem). The homogenate was centrifuged at 13000 rpm for 20 minutes at 4°C. The supernatant was collected and quantified to determine the concentration of proteins in the sample, through the use of Pierce Bicinchoninic Acid Assay (BCA) Protein Assay Kit (Thermo Scientific) method. Proteins were denatured in Laemmli Buffer and boiled for 5 minutes at 98 °C. Proteins were separated by SDS gel-electrophoresis, the membrane was incubated with Milk 5% or BSA 5% for 1 hour at room temperature. Then, the primary antibody of interest was added for 2 hours at RT or overnight at 4°C. After the incubation, the membrane was incubated with specific secondary antibodies for 1 hour at RT. Immunoblots were visualized using SuperSignal West pico/femto Chemioluminescent Substrate (34080-34095, Thermo Scientific) and Chemidoc (Bio-Rad, Hercules, CA, USA). Primary antibodies used were: Rabbit anti-ref(2)P 1:1000, rabbit anti-Atg8a 1:5000 (from Gabor Juhasz, Eotvos Lorand University, Budapest, Hungary), mouse anti-β-tubulin 1:8000 (13-8000, GE Healthcare), mouse anti-Repo 1:20 (8D12, Developmental Studies Hybridoma Bank), rat anti-Elav 1:40 (7E8A10, Developmental Studies Hybridoma Bank), rabbit anti-Mitf 1:200, rabbit anti-phospho-AKT (Ser473) 1:1000 (9271, Cell Signaling), rabbit anti-AKT 1:1000 (9272, Cell Signaling). Secondary antibodies used were rabbit, mouse, rat and chicken HRP-conjugated 1:8000 (Amersham). Protein extraction from NS was performed as in [11]. We used the following primary antibodies: Mouse anti-Vinculin 1:1000 (V4505, Sigma Aldrich), rabbit anti-p-ERK 1/2 (Thr202/Tyr204) 1:1000 (4370, Cell Signaling), rabbit anti-ERK 1/2 1:1000 (4695, Cell Signaling). Western blots protein levels were analyzed using Image Lab (Biorad).

### Brain disaggregation, FACS and sorting analysis

Third instar larval brains were processed as indicated in [47]. After disaggregation, cells were immediately separated using FACS. For sorting analysis, cells were separated using BD FACSDiva 8.0.1, then RNA was extracted from sorted cells as mentioned above (see qPCR analysis).

### Patients’ samples, cell culture and pharmacological treatment

GBM patients’ samples were obtained from Neurosurgery Unit of Fondazione IRCCS Ca’ Granda Ospedale Maggiore Policlinico. GBM samples were processed as previously described [10]. All experiments were performed on 3 V1G1^Low^ and V1G1^High^ patients. NS were treated for 3, 6, 9, 12 and 24 h with BafA1 5 nM and 10 nM (sc-201550, Santa Cruz Biotechnology).

### Electron Microscopy

NS were fixed in 2.5% glutaraldehyde, embedded in 2% agar solution, post-fixed in 1% osmium tetroxide in phosphate buffer, dehydrated and embedded in epoxy resin. Images were captured at 1840X magnification, using a FEI Tecnai G2 20 Transmission Electron Microscope at Alembic – San Raffaele (Milan, Italy).

### Statistical Analysis

All experiments were repeated at least three times for quantification and the mean with standard deviation (S.D.) is shown. P-values are as follows: P* ≤ 0.05; P** ≤ 0.01; P***≤ 0.001. Quantifications were performed with ImageJ and Prism was used for statistical analyses. Sample sizes and statistical methods are detailed in the figure legends.

## Abbreviations

AEL: after eggs laying
AKT: serine/threonine kinase AKT
Atg: autophagy-related
Atg8a: Autophagy related protein 8a
BafA1: bafilomycin A1
CNS: central nervous system
DQ-BSA: DQ-bovine serum albumin
EGFR: Epidermal Growth Factor Receptor
*elav*: *embryonic lethal abnormal vision*
EM: electron microscopy
ERK1/2: extra-cellular signal-regulated kinase 1/2
GBM: glioblastoma
GFP: green fluorescent protein
Lamp1: lysosomal-associated membrane protein 1
MITF: microphthalmia-associated transcription factor
mTOR: mammalian target of rapamycin
mTORC1: the mammalian target of rapamycin complex 1
NS: neurospheres
PI3K: phosphoinositide 3- kinase
ref(2)P: refractory to Sigma P
*repo*: *reverse polarity*
RNAi: RNA interference
S.D.: Standard Deviation
TFEB: the transcription factor EB
TMZ: Temozolomide
V-ATPase: vacuolar-H+-ATPase
ΔhEGFR: constitutively active form of human EGFR

## Acknowledgments

TV acknowledges the support of the UNITECH Nolimits microscopy facility of the university of Milan. This work is supported by the AIRC (Associazione Italiana Ricerca contro il Cancro) Investigator grant 20661 and the WCR (Worldwide Cancer Research) grant 18-0399 to TV and by by Fondazione Cariplo (grant 2014-1148) to VV.

---

**Figure S1.**
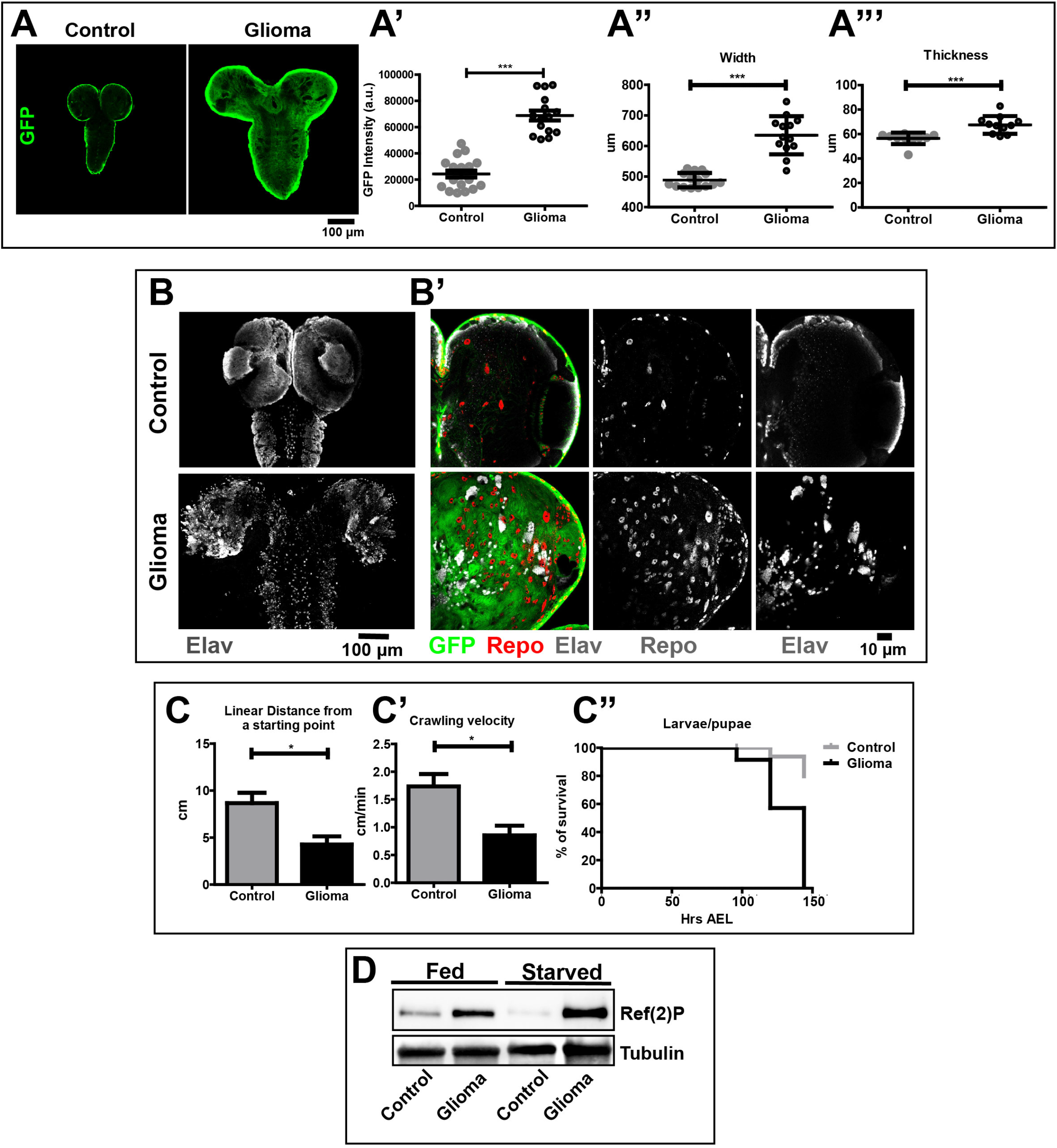
Supporting evidence to Fig. 1. (A-A’) Single medial confocal section of whole CNS of third instar larvae. Dorsal view, anterior up. Larval brains carrying glioma show an increased amount of glia, an enlargement of the CNS and an abnormal ventral nerve cord relative to control brain. Glia quantification is performed taking advantage of glial-specific GFP expression. Notice the increased amount of fluorescence in gliomas compared to controls. n≥10 brains per sample. Data represent the mean ± S.D., *P-value* is determined by Kruskal Wallis test with Dunn’s Multiple Comparison. (A’’-A’’’) The width and thickness (μm) of larval brains are increased in CNS of larvae carrying gliomas compared to controls. n ≥10 brains per sample. The mean ± S.D. are shown, and *P-values* are determined by Mann-Whitney test. (B) Max-projections of medial confocal sections of larval brains. The architecture of Elav cells is strongly compromised in glioma brains compared control brains. (B’) Single medial confocal sections show tissue architecture of *Drosophila* larval brains. Anti-GFP marks glial cell bodies and membranes. These cells cover the entire optic lobe in glioma samples, while mostly the external surface of the optic lobes in control samples. Glial cell nuclei (anti-Repo), are numerically increased in gliomas compared to controls. Neurons, marked by anti-Elav, show a completely subverted morphology in glioma optic lobes compared to control optic lobes, highlighting how the expansion of glia influences neural tissue. Single channels show the nuclei of Repo and Elav cells. (C-C’) The ability to cover a linear distance (cm) and the crawling velocity (cm/min) are both reduced in larvae carrying gliomas. Data represent the mean ± S.D. of n≥3 independent experiments and *P-values* are determined by Mann-Whitney test. (C’’) Survival curves of larvae. Data represent the mean ± S.D. of n≥3 independent experiments and P-value is determined by Log-rank (Mantel-Cox). (D) Western Blot to detect expression of Ref(2)P and Atg8a I-II. Protein levels are evaluated in fed and starved conditions. Starvation induces ref(2)P degradation in control animals (lane 3 vs 1), while further increase in glioma (lane 4 vs 2). Tubulin is used as a loading control.

**Figure S2.**
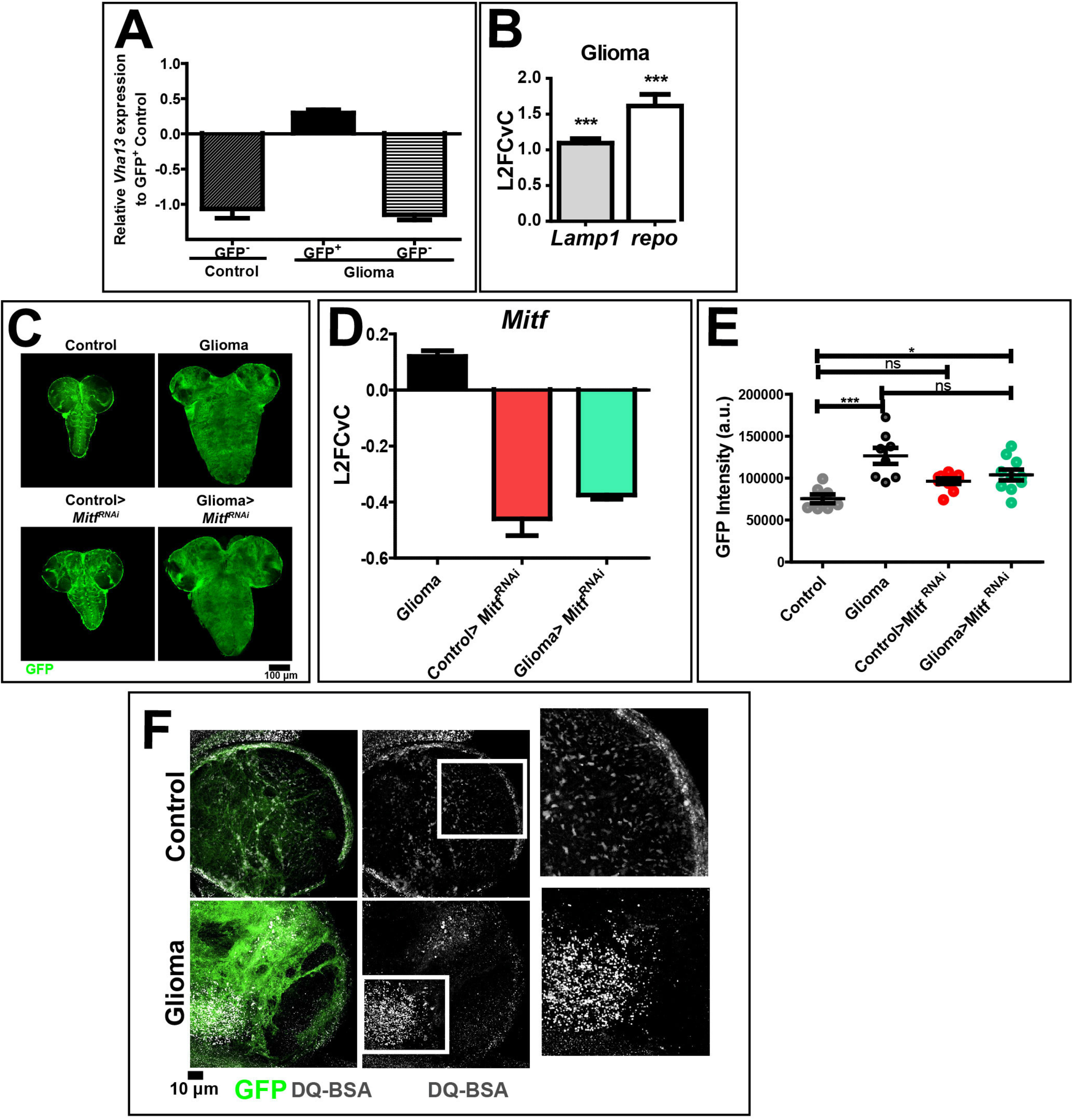
Supporting evidence to Fig. 2. (A) Expression of *Vha13* after tissue disaggregation GFP^−^ or GFP^+^ glial cells that belong to control and glioma. Normalization is relative to GFP+ cells of control brains. Data represent the mean ± S.D. of n≥3 independent experiments. (B) *Lamp1* and *repo* mRNA expression in glioma CNSs. Data represent the mean ± S.D. of n≥3 independent experiments, Mann-Whitney test. (C) Single medial confocal section of whole CNS of third instar larvae. Dorsal view, anterior up. Larval brains carrying gliomas show in which *Mitf* has been downregulated show similar growth to controls. (D) *Mitf* mRNA expression in extracts of the indicated genotypes evaluated by qPCR. Samples carrying the downregulation of *Mitf* show a markedly reduced expression of the gene. Notice that mRNA levels of *Mitf* are upregulated in gliomas compared to controls. Data represent the mean ± S.D. of n≥2 independent experiments. (E) Quantification of glial cells by immunostaining using anti-GFP. Downregulation of *Mitf* in gliomas slightly decreases glial overgrowth. Mean ± SEM are shown, *P*-*values* are determined by Kruskal Wallis test with Dunn’s Multiple Comparison. (F) Max projections of medial confocal sections of third instar larval CNSs, after 6 hrs of incubation with DQ-BSA. Anti-GFP stains glial cell membranes, while DQ-BSA measures protein degradation in lysosomes. Glioma brains preserve their lysosomal activity. High magnifications of insets show the lysosomal expansion in gliomas.

**Figure S3.**
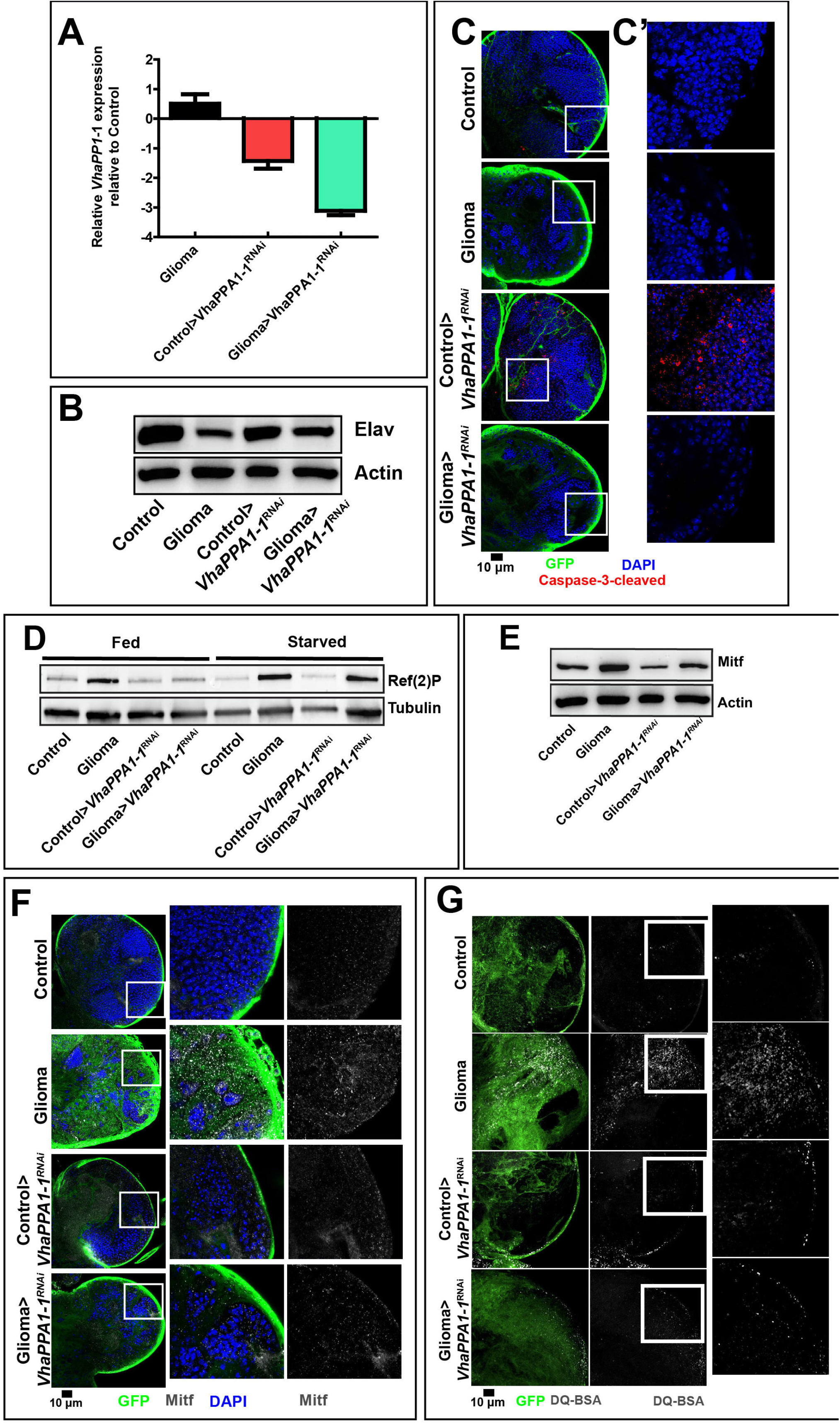
Supporting evidence to Fig. 3. (A) *VhaPPA1-1* expression evaluated by qPCR. Samples carrying the downregulation show reduced expression of the gene. Data represent the mean ± S.D. of n≥3 independent experiments. (B) Western blot showing levels of Elav in the indicated samples. Actin is used as a loading control. n≥3 independent experiments. (C) Single medial confocal sections of *Drosophila* larval CNSs. Cell nuclei are stained with DAPI, glial cell membranes are stained with anti-GFP, apoptotic cells are stained with anti-cleaved caspase-3. High magnifications of insets show apoptosis in control>*VhaPPA1-1*^*RNAi*^. (D-E) Western blot showing levels of Ref(2)P (D) or Mitf (E) in the indicated samples conditions. Tubulin (D) o Actin (E) are used as a loading control. (F) Single medial confocal sections of third instar larval CNSs stained as indicated. High magnifications insets show Mitf subcellular localization, which is almost exclusively cytoplasmic in all samples. (G) Max projections of medial confocal sections of third instar larval brains after 6 hrs of incubation with DQ-BSA.

## Supplement methods

### Immunofluorescence

Mouse anti-Repo 1:20 (Developmental Studies Hybridoma Bank, 8D12), Rat anti-Elav 1:40 (Developmental Studies Hybridoma Bank, 7E8A10), Rabbit anti-Cleaved Caspase 3 1:200 (Cell Signaling, 9661), Mouse-anti Actin 1:500 (Sigma)

### qPCR analysis

*Lamp1 F_5’GCTTTCCTTTATGCAAATTCATC3’, Lamp1 R_5’GCTGAACCGTTTGATTTTCC3’; Rpl32 F_5’CGGATCGATATGCTAAGCTGT3’ Rpl32 R_ 5’CGACGCACTCTGTTGTCG3’;*

### DQ Red BSA assay

DQ Red BSA (Thermo Fisher) was used at a concentration of 120μg/μl diluted in M3 insect medium (S3652, Sigma). Samples were incubated for 6 hours at room temperature, then were fixed with 4% PFA. Brains were mounted on a slide with glycerol 70% for the subsequent confocal acquisitions.

### Locomotor Assay

Using a brush, we moved third instar larvae to a lid of a 15 cm Petri dish (pre-coated with 2% agarose). Under the lid is pasted a graph paper with a 0.2 cm^2^ grid. We measured the number of the grid lines crossed in 5 minutes. This value is the linear distance from a starting point, while the crawling velocity is obtained dividing the previous value for the time.

### Survival Assay

Mated flies released eggs for 3 hours on new vials. Eggs were grown at 25 C° and checked every day until 150 hrs after eggs laying (AEL).

---

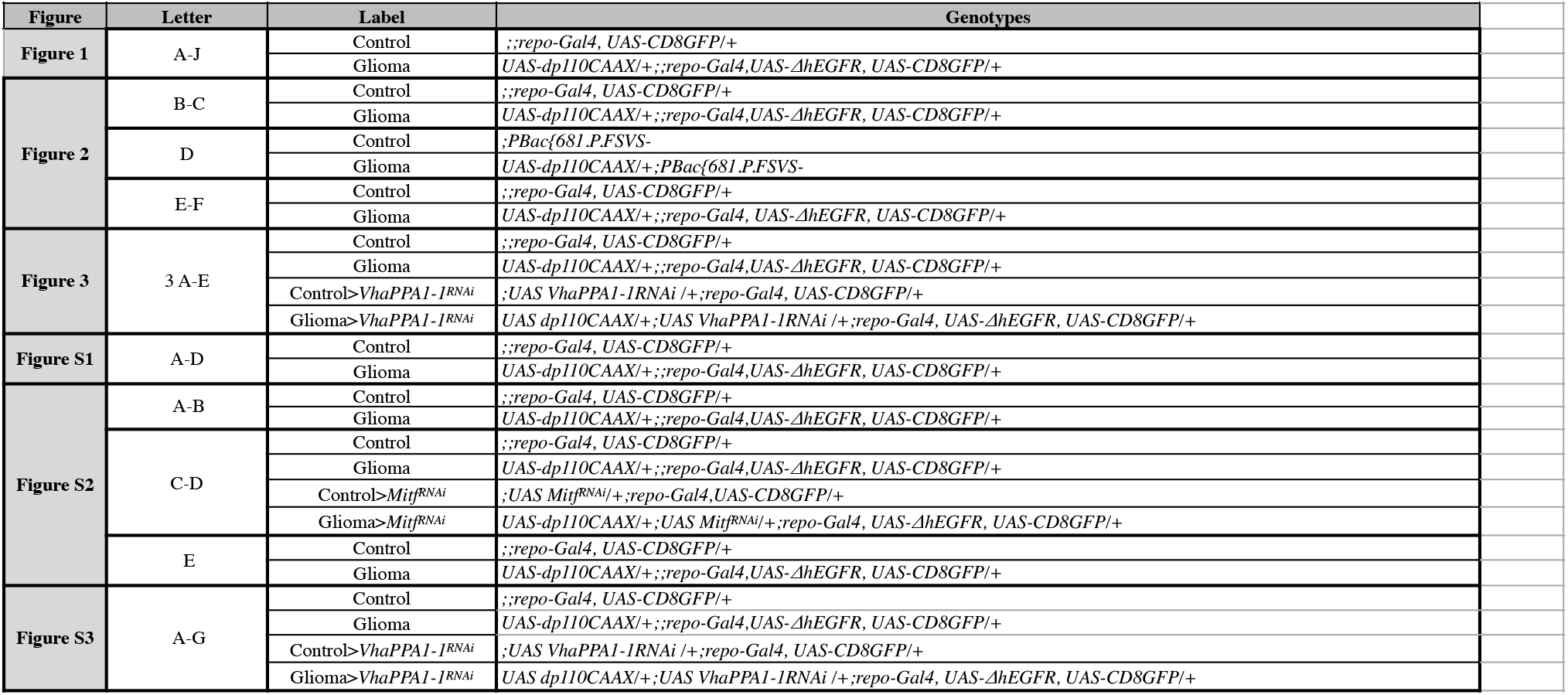

## Notes

### Competing Interest Statement

The authors have declared no competing interest.

